# Cyclic nucleotide-induced superhelical structure activates a bacterial TIR immune effector

**DOI:** 10.1101/2022.05.04.490601

**Authors:** Gaëlle Hogrel, Abbie Guild, Shirley Graham, Hannah Rickman, Sabine Grüschow, Quentin Bertrand, Laura Spagnolo, Malcolm F White

## Abstract

Cyclic nucleotide signalling is a key component of anti-viral defence in all domains of life, from bacteria to humans. Viral detection activates a nucleotide cyclase to generate a second messenger, resulting in activation of effector proteins. This is exemplified by the metazoan cGAS-STING innate immunity pathway ^1^, which originated in bacteria ^2^. These defence systems require a sensor domain such as STING or SAVED to bind the cyclic nucleotide, coupled with an effector domain that causes cell death when activated by destroying essential biomolecules ^3^. One example is the TIR (Toll/interleukin-1 receptor) domain, which degrades the essential cofactor NAD^+^ when activated in response to pathogen invasion in plants and bacteria ^2,4,5^ or during nerve cell programmed death ^6^. Here, we show that a bacterial anti-viral defence system generates a cyclic tri-adenylate (cA_3_) signal which binds to a TIR-SAVED effector, acting as the “glue” to allow assembly of an extended superhelical solenoid structure. Adjacent TIR subunits interact to organise and complete a composite active site, allowing NAD^+^ degradation. Our study illuminates a striking example of large-scale molecular assembly controlled by cyclic nucleotides and reveals key details of the mechanism of TIR enzyme activation.

---

Recent discoveries have revealed a central role for cyclic nucleotide second messengers in prokaryotic anti-viral defence by type III CRISPR ^7,8^, CBASS (Cyclic nucleotide Based Antiphage Signalling Systems) ^3^ and PYCSAR (Pyrimidine Cyclase System for Antiphage Resistance) ^9^. These systems activate potent effector proteins that destroy key cellular components such as nucleic acids, cofactors or membranes to disrupt viral replication ^3,10^. A good example is the TIR domain, which functions as an enzyme that degrades NAD^+^ to cause cell death in plants infected with pathogens ^5,11^, anti-phage immunity in the bacterial Thoeris ^12,13^ and TIR-STING ^2^ systems, and programmed nerve cell death in metazoa ^6^. TIR domain activation requires effector subunit assembly, but the molecular mechanisms are still not fully understood.

## The TIR-SAVED effector is activated by cA_3_ to degrade NAD^+^

Type III CRISPR and CBASS systems both use the SAVED (SMODS-associated and fused to various effector domains) cyclic nucleotide binding sensor domain ^14,15^. Here we focussed on a Type II-C CBASS from the Gram-positive bacterium *Microbacterium ketosireducens* (Mke) ^16^ which has a nucleotide cyclase (CD-NTase), TIR-SAVED and NucC effectors, along with an ubiquitin-like modification system of unknown function (Fig. 1a). We designed synthetic genes for the expression of MkeCD-NTase and MkeTIR-SAVED in *E. coli* and purified the recombinant proteins using cleavable N-terminal his-tags and gel filtration (Extended Data Fig.1). We investigated the activity of the cyclase by incubating the protein with a range of nucleotides, including α-^32^P-ATP for visualisation, and analysis by thin layer chromatography (TLC). A radioactive product running at the position of a cA_3_ standard was observed when ATP was present in the reaction (Fig. 1b), and the addition of other nucleotides did not result in any observable change (Extended Data Fig.2a). The analysis of the reaction products by liquid chromatography confirmed that the cyclase uses ATP to produce cA_3_ that co-elutes with a synthetic 3’,3’,3’-cA_3_ standard (Fig. 1c).

**Fig. 1.**
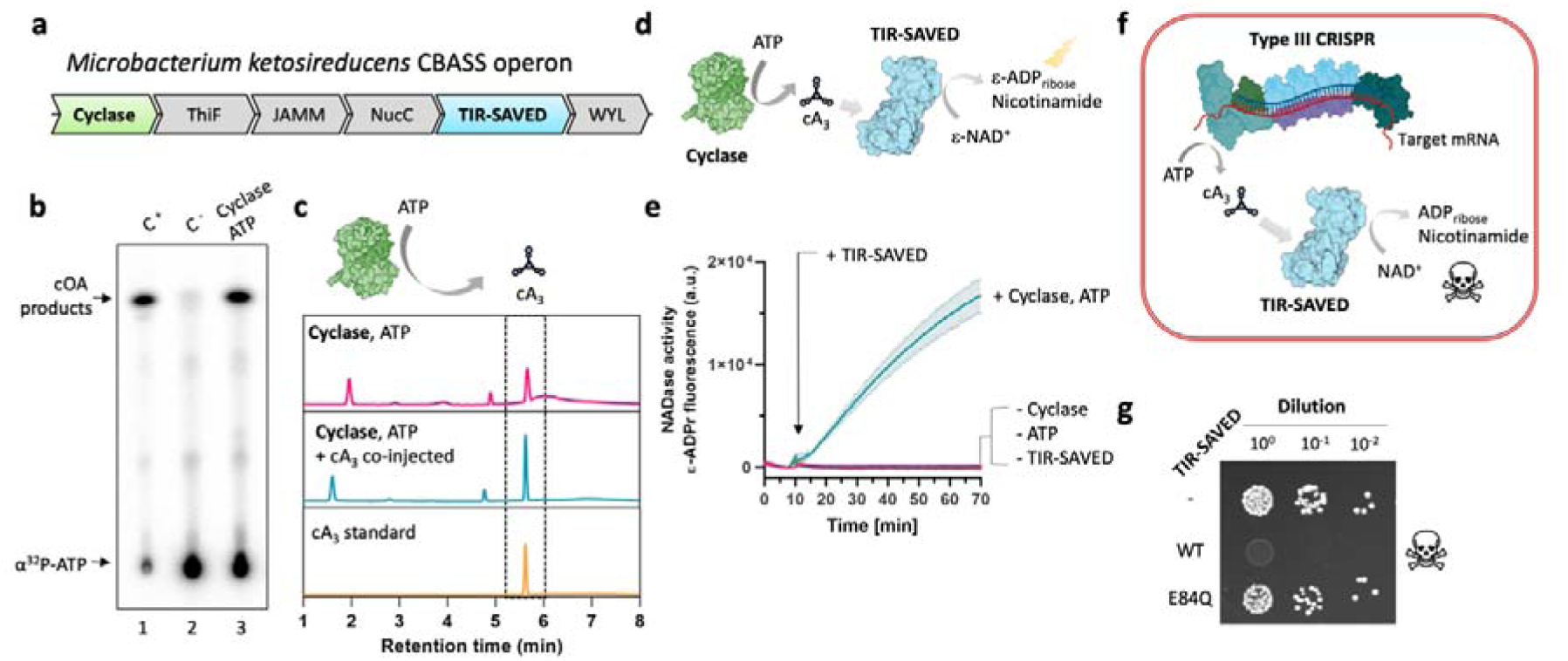
MkeCBASS generates a cA_3_ second messenger to activate the TIR-SAVED NADase effector. **A**, MkeCBASS operon (RS81_contig000028). **b**, The cyclase (CD-NTase) uses ATP to produce cyclic-oligoadenylate (cOA) products (lane 3), visualised after TLC. Negative control without protein (lane 2), positive control (lane 1) cA_3_ produced using the Type III CRISPR complex of *Vibrio metoecus* ^21^. **c**, HPLC analysis of reaction products confirmed the cA_3_ product. Raw figure available in Supplemental Data Fig.1. **d**, Schematic of the reaction used to analyse TIR-SAVED NADase activity. **e**, Cyclase products activate TIR-SAVED NADase activity. The CD-NTase was incubated with ATP for 2 h before adding ε-NAD^+^ and TIR-SAVED. Data are means plotted with standard deviation for triplicate experiments. **f**, Schematic of the plasmid challenge assay. The Type III CRISPR complex produces cA_3_ on binding the target transcript, activating TIR-SAVED NADase activity. **g**, *E. coli* transformants after plasmid immunity assay of *Mtb*Csm (Type III CRISPR system) combined with TIR-SAVED. Wild-type TIR-SAVED prevents the growth of transformants whereas the inactive E84Q variant does not. Additional conditions, replicates and details of the constructs are shown in Extended data Fig.3.

To investigate the activity of the TIR-SAVED effector, we used □-NAD^+^, an NAD analogue that emits a fluorescent signal on cleavage by TIR proteins (Fig. 1d) ^2,11^. We screened a range of commercially available cyclic nucleotide molecules for the ability to activate the TIR-SAVED effector, observing that only cA_3_ resulted in generation of a fluorescent signal (Extended Data Fig. 2b). We proceeded to couple the cyclic nucleotide production by the cyclase with the NADase assay to follow the activation of TIR-SAVED. In the presence of ATP and a high pH reaction buffer, the cyclase activated the TIR-SAVED effector to degrade NAD^+^ (Fig. 1e). Together these data demonstrate that the MkeCD-NTase synthesises a 3’,3’,3’-cA_3_ product that can activate the TIR-SAVED NADase activity.

The initial rate of NAD^+^ cleavage increased linearly with cA_3_ concentration up to a value of 0.5 µM cA_3_, which corresponded with the concentration of TIR-SAVED in the assay, consistent with a 1:1 molar ratio of cA_3_ to TIR-SAVED in the activated form of the effector (Extended Data Fig. 2c). We next examined the Michaelis-Menten parameters of TIR-SAVED NADase activity, determining a K_M_ of 470 µM and a k_cat_/K_M_ of 2.08 × 10^3^ M^-1^s^-1^ (Extended Data Fig. 2d). This K_M_ value fits in the concentration range of NAD^+^ found in mammalian cells and *E. coli* (200-640 µM) ^17,18^. Various bacterial TIR proteins have estimated K_M_ values between 196 and 488 µM ^4^. In the *Thoeris* system of *Bacillus cereus*, the NADase enzyme ThsA activated by cyclic ADP ribose (cADPr) has a K_M_ of 270 µM for NAD and a catalytic efficiency (k_cat_/K_M_) of 2.1 × 10^3^ M^-1^s^-1 13^, in good agreement with our observations.

## Activation of the TIR-SAVED effector in *E. coli* leads to cell death

CBASS defence systems tend to be phage and host species specific, and the mechanism of activation in response to phage infection remains largely unknown ^19^. As we could not analyse CBASS activity in the cognate host, we took advantage of the observation that TIR-SAVED is activated by cA_3_ by coupling the effector with a type III CRISPR system from *Mycobacterium tuberculosis (Mtb)*, which generates a range of cyclic oligoadenylate species including cA_3_ on detection of target RNA ^20^. Here, we replaced the cognate Csm6 effector with MkeTIR-SAVED and programmed the CRISPR system with a guide RNA complementary to the tetracycline resistance gene *tetR* (Fig. 1f, Extended Data Fig.3a). When the active CRISPR system was present along with wild-type TIR-SAVED, transformation of a target plasmid containing the *tetR* gene resulted in no cell growth on plates containing tetracycline (Fig. 1g). This phenotype required the production of cA_3_ (Extended Data Fig.3) and the NADase activity of TIR-SAVED, as variant E84Q, which targets the active site (Fig. 2), relieved the effect ^2,11,12^. This suggests that TIR-SAVED, activated by cA_3_, is responsible for cell death by NAD^+^ hydrolysis.

**Fig. 2.**
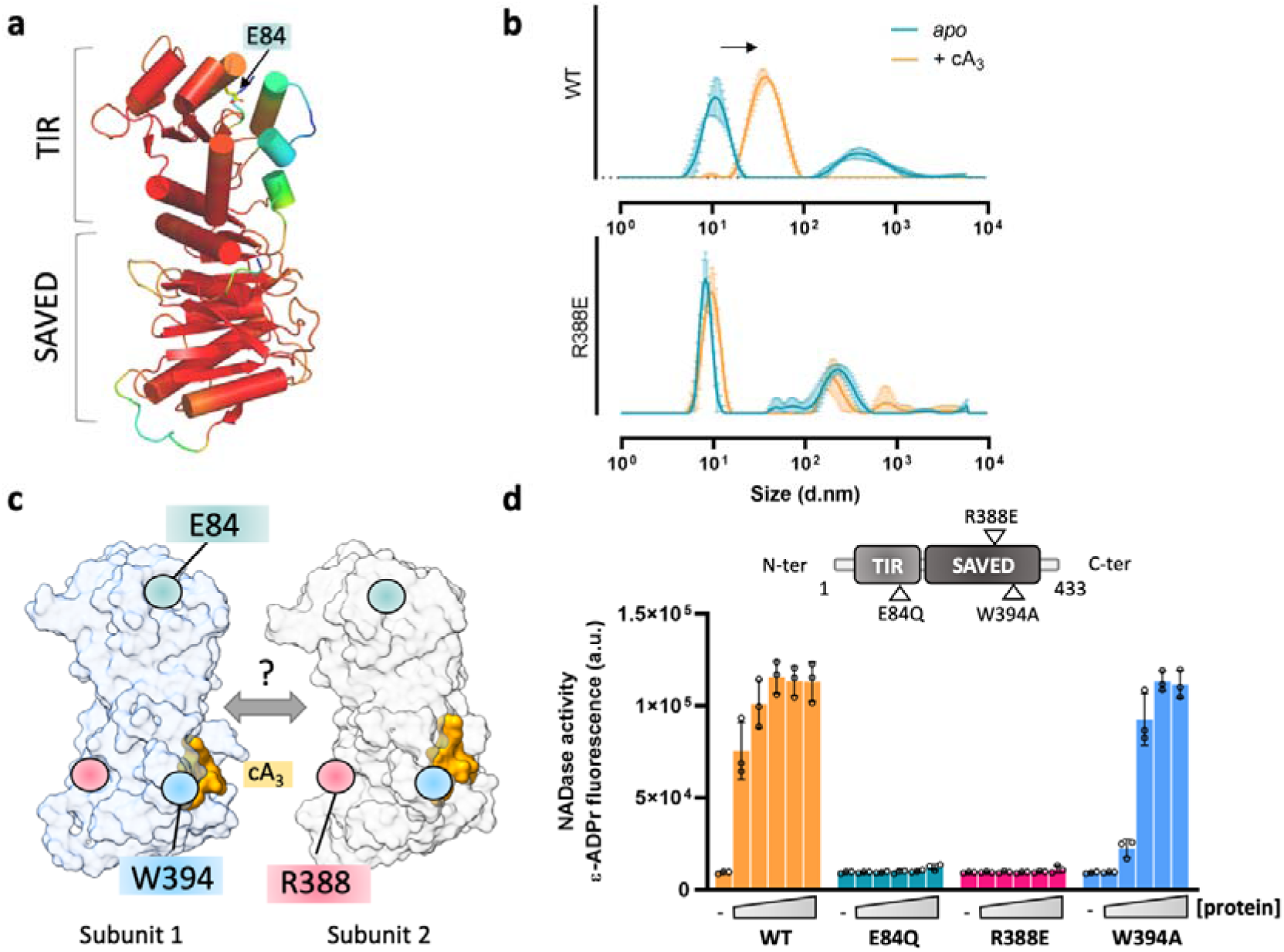
cA_3_ drives TIR-SAVED oligomerisation important for NADase activity. **a**, Structural prediction of TIR-SAVED generated using AF2, coloured based on IDDT score (red (high) to blue (low)) obtained for this model. **b**, Dynamic light scattering shows that the wild-type, but not the R388E variant of TIR-SAVED increases in size on cA_3_ binding. **c**, Head-to-tail association model of two TIR-SAVED subunits. Key residues used for mutagenesis are highlighted. **d**, NADase activity of TIR-SAVED variants. A range of protein concentration (0, 0.16, 0.5, 1.5, 4, 13.5 µM) was incubated with 27 µM cA_3_ and 500 µM □-NAD^+^ for 60 min. Data are means plotted with standard deviations for triplicate experiments.

## TIR-SAVED is a monomer and multimerises on addition of cA_3_

The structure of the TIR-SAVED monomer (Fig. 2a) was predicted using Alphafold2 (AF2) ^22^ as implemented by the Colabfold server ^23^. Five models were predicted, which were essentially superimposable with IDDT values ranging from 91.1 to 90.1. Regions of the protein predicted with less certainty included the N- and C-termini and loops predicted at positions 40-51, 175-183 and 285-294 (more details in Supplemental data Fig.2). Comparison of the AF2 structure with the structure of the Cap4 protein from *Acinetobacter baumannii* ^24^ (PDB code 6WAN) gave a Z-score of 14.1 and an RSMD of 3.9 Å for the SAVED domain (sequence alignment in Supplemental Data Fig. 3). The two domains are shown overlaid in Extended data Fig.4, allowing the positioning of the cA_3_ activator in the conserved binding site. Notably, W394 is conserved in both proteins and predicted to form an extensive interaction with one of the adenine bases. The modelled TIR domain is most similar to other catalytic TIR domains such as those found in the plant NLR RPP1 immune receptor complex (PDB code 7CRC; Z-score of 13.6 and RMSD of 4.1 Å over 183 aa) ^25^.

Analytical size exclusion chromatography (SEC) combined with Small Angle X-ray scattering (SAXS) indicated that TIR-SAVED tends to be monomeric in solution (Extended Data Fig. 5a). This result was also supported by analytical gel filtration (Supplemental Data Fig. 4). The theoretical scattering curve obtained for the AF2 model fitted the SAXS profile of TIR-SAVED in solution with a χ^2^ 1.139 (Extended Data Fig.5b). Addition of cA_3_ to the protein (1.5:1 molar ratio), induced a dramatic shift in the elution volume on gel filtration, with the monomeric species replaced by a broad peak of absorbance eluting much earlier, suggesting the presence of high molecular weight complexes (Extended data Fig. 5c). We confirmed this by Dynamic Light Scattering (DLS), which showed that the global particle size was significantly increased upon cA_3_ addition (Fig.2b). These data strongly suggest that cA_3_ binding induces multimerization of the TIR-SAVED protein, reminiscent of that observed for the Cap4 protein from several species, where head-to-tail multimers of two or three subunits were observed by electron microscopy ^24^. We therefore investigated the possibility of a head-to-tail dimer of TIR-SAVED sandwiching a single cA_3_ molecule, where subunit 1 binds cA_3_ in the canonical binding site and subunit 2 provides a second interface on the opposite side of the protein (Fig. 2c). To confirm the prediction of the canonical binding site, we analysed the W394A variant, which showed weaker cA_3_ binding (Extended Data Fig.6a), and a reduction of cA_3_-activated NADase activity *in vitro* (Fig. 2d, Extended Data Fig. 6b).

The model also allowed us to predict a potential role for a conserved arginine residue, R388 (see sequence alignment in Extended Data Fig. 7), in subunit 2, which is suitably positioned to act as a ligand for the negatively charged cyclic nucleotide ligand (Fig. 2c). The R388E variant of TIR-SAVED was unable to multimerize as assessed by DLS (Fig. 2b, lower panel), but could still bind cA_3_ as observed in electrophoretic mobility shift and thermal shift assays (Extended Data Fig. 6), consistent with cA_3_ binding only at the canonical subunit 1 site in the monomer. Furthermore, the R388E variant was no longer capable of being activated by cA_3_ to function as an NADase (Fig. 2d). Together, these data suggest that the TIR-SAVED protein multimerises in a head-to-tail configuration on binding cA_3_, and that this structural change is linked to the activation of the TIR enzymatic domain.

### Structure of the active TIR-SAVED complex

Cryo-electron microscopy demonstrated that cA_3_ drove the formation of ordered TIR-SAVED filaments (Fig. 3a). The cryo images showed the co-existence of both side views and top views. The assembly is characterised by a right-handed superhelical solenoid with 22 nm radius and a 14 nm pitch. 17 TIR-SAVED monomers are present in each turn of the filament (Fig. 3a). Local resolution analysis performed with the CryoSPARC package highlighted a slightly anisotropic resolution, going from 2.5 to 5 Å (Extended data, Figure 8). 3D variability analysis of a structure including two pitches of the assembly highlighted a degree of flexibility over the filament (Supplemental Data Fig.5 and Fig.6), as shown in and the supplementary videos. These show a slight variability in the radius of the filaments, as well as in its pitch, therefore we chose to proceed with the analysis of one tier of the complex in isolation.

**Fig. 3.**
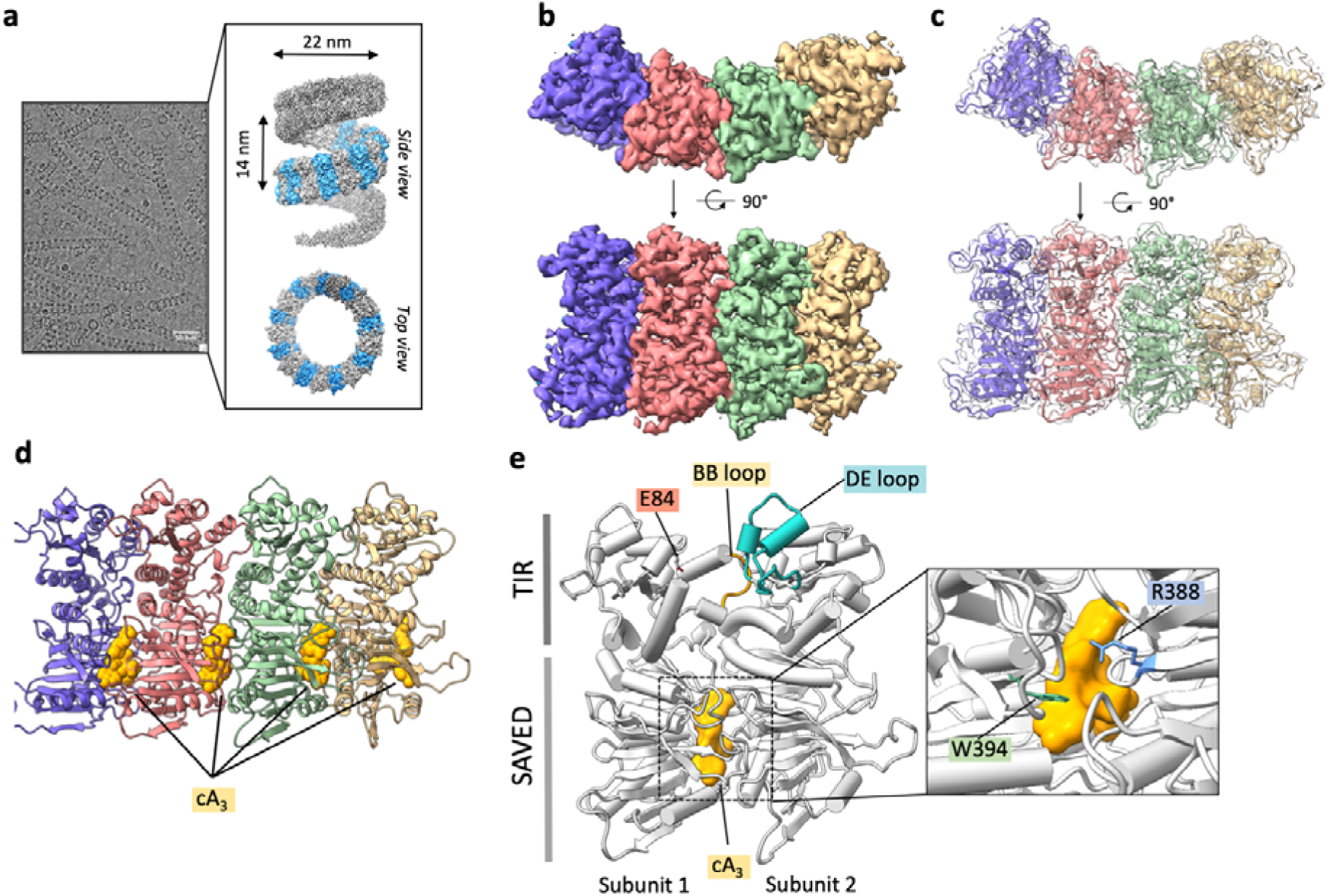
Structure of the activated TIR-SAVED/cA_3_ assembly. **a**, Cryo-EM micrograph of TIR-SAVED in presence of cA_3_ (1:1.5 molar ratio) (left panel), TIR-SAVED/cA_3_ filament density map (right panel). **b**, Cryo-EM density of the final processed tetramer. Orientation with the TIR domain on the top. **c**, Atomic model fitted into the cryo-EM density (PDB: 7QQK). **d**, Final atomic model of TIR-SAVED with cA_3_ packed between each of the four subunits. **e**, Analysis of TIR-SAVED/cA_3_/TIR-SAVED interaction interfaces based on the cryo-EM model.Key residues are highlighted.

To build a model of TIR-SAVED, we used four copies of the AF2 output as a starting model for rigid body fitting using the Chimera program. The rigid-body fitted tetramer was then refined as described in the methods. Four densities that correspond to the characteristic shape and size of the cA_3_ ligand ^24^, visible in the outstanding density, were fitted and refined. The final atomic model here confirms the head-to-tail assembly where the cA_3_ is located on the previously described binding pocket of SAVED1 (head subunit), next to W394, and closed by the back side of SAVED2 (tail subunit) (Fig. 3e). The cA_3_ is tightly bound between SAVED1-SAVED2 with contact residues from both subunits (Extended Data Fig. 9). The position of R388 in SAVED2 at the interface is consistent with the phenotype of the R388E variant.

We next wished to understand whether the assembly of the helical structure was essential for TIR-SAVED activation. To investigate this, we combined two variant proteins, E84Q and R388E, that are completely inactive in isolation as the former can multimerise but is catalytically dead whilst the latter has a wild-type active site but cannot multimerise on its own (Fig. 4a). By combining the two variants at a 1:1 ratio, which allows the formation of dimers with one intact composite cA_3_ binding site and one active site, we observed the recovery of NADase activity in the presence of cA_3_ (Fig. 4b). This demonstrates that the dimeric TIR-SAVED structure is sufficient to trigger TIR activity. As the concentration of the catalytically dead E84Q variant in the mixture was increased to ratio of 2:1 with respect to R388E, we observed a progressive increase in NADase activity, consistent with a role for the extended helical structure for full activation of the TIR domain (Fig. 4b).

**Fig. 4.**
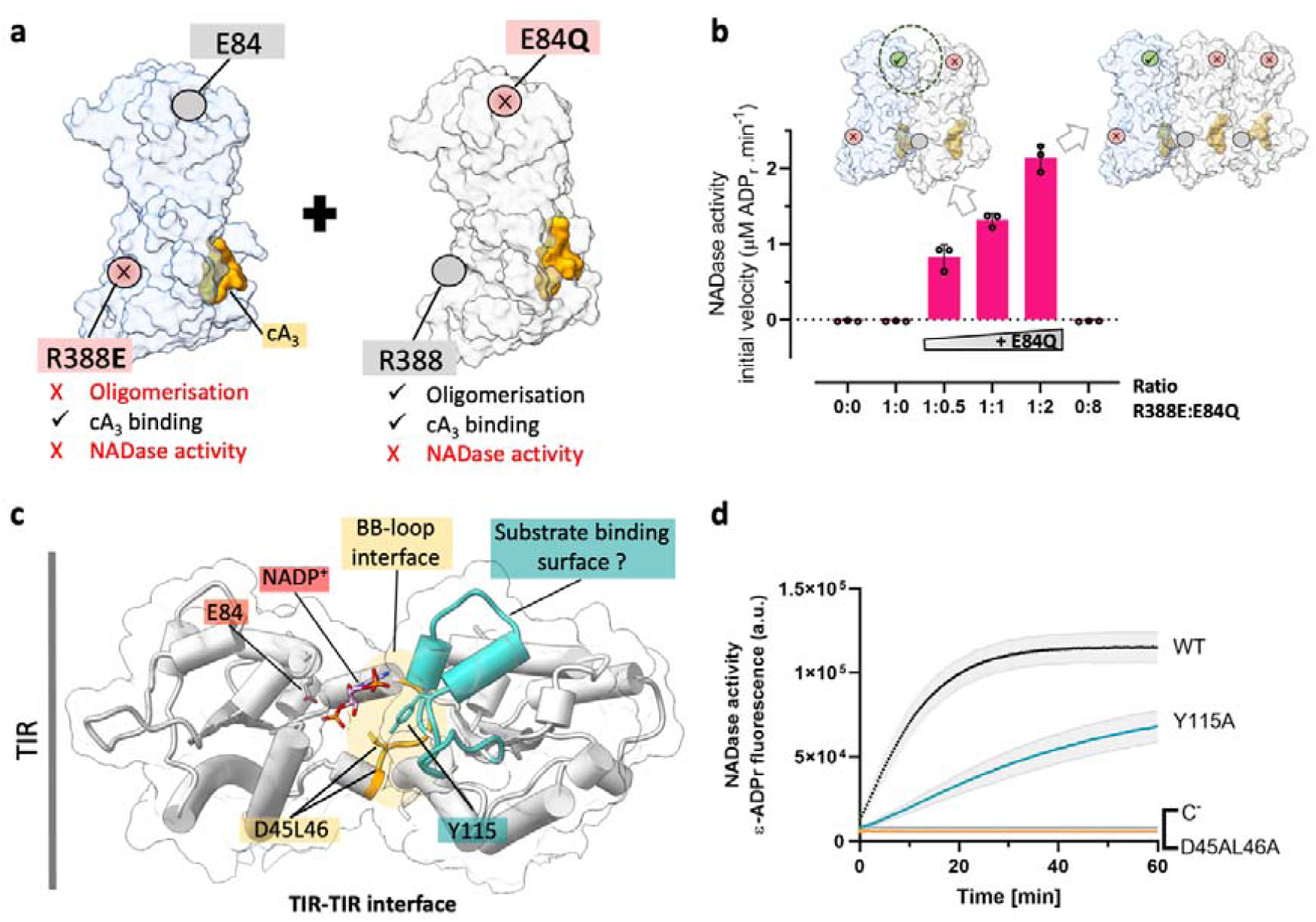
Multimerisation of TIR-SAVED generates a composite TIR active site. **a**, Schematic and properties of the two mutants R388E and E84Q combined for NADase assays. **b**, Initial NADase activity of 0.25 µM R388E in presence of increased E84Q concentration incubated with 500 µM □NAD^+^ and 2 µM cA_3_. Experiments in triplicate with standard deviation shown. **c**, MkeTIR dimer interface (cryo-EM model) with NADP+ modelled inside based on structural alignment with *Vitis rotundifolia* RUN1 TIR domain (PDB: 6o0W). **d**, NADase activity of D45AL46A and Y115A variants compared to WT (1.5 µM TIR-SAVED incubated with 500 µM □NAD^+^ in presence of 27 µM cA_3_). The mean of triplicate experiments are shown with standard deviation for the WT and Y115A variant; the inactive D45AL46A was assayed in duplicate.

## Mechanism of TIR enzyme activation

TIR domains are ubiquitous, performing both protein scaffolding and enzymatic roles in different contexts across all three domains of life (reviewed in ^4^). As is the case for TIR-SAVED, the catalytically active TIR domains, which degrade NAD^+^ to cause cell death, typically rely on multimerization linked to activation ^2,4,5,11,25,26^, but the molecular mechanism for this activation is not well understood. In the filament assembly, TIR domains display a conserved interaction interface involving the BB loop of subunit 1, which appears to be held in an open configuration due to interaction with the DE loop from the adjacent subunit (Fig. 3e). This BB-loop interface is also observed in TIR proto-filament formation such as for the catalytically inactive TIR domain of the human MAL Toll-like receptor protein ^27^ and also the Run1-TIR tetramer. The BB loop is suspected, in other NAD^+^ consuming TIR proteins, to regulate the access to the active site ^11,28 25,26^ and it has been proposed recently for the RUN1-TIR domain that the DE loop of the adjacent subunit contributes to NAD binding ^28^.

We therefore tested the hypothesis that a composite NADase active site is formed at the interface between two adjacent TIR domains in the TIR-SAVED filament (Fig. 4c). We first investigated the BB loop, showing that a variant protein with the double mutation D45A/L46A completely lacked NADase activity (Fig. 4d) without affecting cA_3_-dependent multimerization (Extended Data Fig. 10). We proceeded to explore the role of putative NAD^+^ binding site by mutating the highly conserved residue Tyr-115, which is suitably positioned to interact with NAD^+^ in the TIR-SAVED filament (Fig. 4c). The Y115A variant had only 10% of the NADase activity of the wild-type enzyme, together with a 3-fold increase in K_M_ for εNAD (Fig. 4d, Extended Data Fig. 10a). This supports the model of a composite NADase active site that has also been proposed for the TIR-NLR RPP1 immune receptor ^25^ and which may therefore have broad relevance for catalytic TIR proteins.

## Concluding Remarks

Here we have explored a CBASS system with a TIR-SAVED effector that is activated by cA_3_. TIR-SAVED proteins have not previously been studied, but their constituent domains are found in a number of other contexts. The SAVED domain is a distant cousin of the cOA-sensing CARF domain associated with type III CRISPR systems ^15^, and is found in 30% of CBASS operons ^24^. SAVED domains fused to nucleases have been shown to bind cyclic di-and tri-nucleotides, activating the associated nuclease domain for DNA degradation ^24,29^. Our data suggest that head-to-tail stacking of SAVED domains, potentiated by cyclic nucleotide binding, is a defining feature of effector activation. Oligomerisation with “open symmetry” seems to be a widespread property of prokaryotic innate immune effectors, encompassing the bacterial STING, SAVED and PYCSAR proteins ^2,9,24^. We have demonstrated that this oligomerisation is essential for TIR-SAVED function; firstly, as it allows access to the active site by prising open the BB loop; secondly, as residues from the adjacent subunit participate in the catalytic cycle. Thus, the bacterial TIR-domain effectors appear to conform to the emerging paradigm of “signaling by cooperative assembly formation” (SCAF) proposed for the eukaryotic signalling complexes ^30^. Indeed, oligomerisation-dependent activation of bacterial TIR effectors is likely to be the ancestral mechanism underlying the whole family of catalytic and non-catalytic TIR signalling complexes. This is exemplified by the accompanying study of the bacterial TIR-STING effector ^31^. It remains to be determined how large the TIR-SAVED helices grow *in vivo* in bacteria; this is likely to be influenced by the copy number of the effector protein in cells, and on the concentration of cyclic nucleotide generated by activation of the cyclase enzyme. Our subunit mixing experiments demonstrate that although the minimal head-to-tail dimer has detectable NADase activity, formation of short helical segments increases this significantly. There is also the question of whether these helices can disassemble, for example due to degradation of the cyclic nucleotide “glue” by a ring nuclease ^10^. These are fascinating areas for future studies.

## Acknowledgements

This work was funded by the BBSRC (ref. BB/S000313 and BB/T004789) and a European Research Council Advanced Grant (grant number 101018608) to MFW. We thank Dr Januka Athukoralage, Dr Tracey Gloster and Mr Stuart McQuarrie for discussions. We thank M. Tully for assistance in using beamline BM29. We acknowledge the Scottish Centre for Macromolecular Imaging (SCMI), Dr Mairi Clarke and Dr James Streetley for assistance with cryo-EM experiments and access to instrumentation, funded by the MRC (MC_PC_17135) and SFC (H17007). This work used the platforms of the Grenoble Instruct-ERIC center (ISBG; UAR 3518 CNRS-CEA-UGA-EMBL) within the Grenoble Partnership for Structural Biology (PSB), supported by FRISBI (ANR-10-INBS-0005-02) and GRAL, financed within the University Grenoble Alpes graduate school (Ecoles Universitaires de Recherche) CBH-EUR-GS (ANR-17-EURE-0003).

## Author Contributions

G.H. planned, carried out and analysed the biochemical experiments and structural analyses and drafted the manuscript; A.G. collected and analysed Electron Microscopy data; S. G. cloned, expressed and purified the wild-type and variant proteins; H.R. carried out the preliminary biochemical analysis of the effector protein; S.Gu analysed the cyclic nucleotides; Q.B. carried out the SAXS; L.S. planned and analysed the EM analyses; M.F.W. conceptualised and oversaw the project, obtained funding and analysed data along with the other authors. All authors contributed to the drafting and revision of the manuscript.

## Additional information

Supplementary information is available for this paper.

Correspondence and requests for materials should be addressed to Malcolm White or Laura Spagnolo.

## Methods

### Cloning & mutagenesis

The synthetic genes encoding *Microbacterium ketosireducens* (Mke) *cdn* and *tir-saved* were codon optimised for expression in *Escherichia coli* and purchased from Integrated DNA Technologies (IDT), Coralville, USA. Genes were cloned into the pEhisV5Tev vector ^32^ between the NcoI and BamHI restriction sites. The constructs were transformed into competent DH5α (*E. coli*) cells and plasmids were extracted using GeneJET plasmid miniprep kit (Thermo Scientific) to verify sequence integrity by sequencing (Eurofins Genomics). Then plasmids were transformed into *E. coli* C43 (DE3) cells for protein expression. MkeTIR-SAVED variants, E84Q, R388E, W394A, D45AL46A and Y115A were generated on both constructs containing *mketir-saved* gene: (i) the expression vector (pEhisV5spacerTev) and (ii) the vector used for plasmid immunity assay (pRAT-Duet).

### Protein expression and purification

Recombinant MkeCD-NTase and MkeTIR-SAVED wild-type and variants were overexpressed in *E. coli* C43 (DE3) cells. After growing at 37°C in LB medium until OD_600_ reached 0.6-0.8, cultures were induced with 0.4 µM final IPTG concentration and incubated at 16°C overnight. Cells were collected by centrifugation and pellets stored at -80°C. Cell pellets were lysed by sonication in Buffer A (50 mM Tris-HCl, pH 7.5, 0.5 M NaCl, 10 mM imidazole, 10% glycerol) supplemented by 1 mg.mL^-1^ final lysozyme concentration and protease inhibitors (cOmplete, EDTA-free protease inhibitor cocktail, Roche). Clarified lysates were loaded onto a 5 mL HiTrap FF column (GE Healthcare), pre-loaded with Ni^2+^ and equilibrated with Buffer B (50 mM Tris-HCl, pH 7.5, 0.5 M NaCl, 30 mM imidazole, 10% glycerol). Then histidine-tagged proteins were eluted along a linear gradient (50 mM to 500 mM imidazole). Recombinant histidine-tagged TEV protease was incubated with the eluted proteins to cleave the histidine-tag and the overall was dialysed overnight at room temperature against Buffer B. Dialysed samples were loaded again onto the Ni^2+^/loaded 5 mL HiTrap FF column (GE Healthcare) equilibrated with Buffer B. The histidine-tagged cleaved proteins where then loaded onto a 26/60 Superdex 200 size-exclusion column (GE Healthcare) equilibrated with buffer C: 20 mM Tris-HCl, pH 7.5, 0.25 M NaCl, 10% glycerol. Proteins were concentrated with 30-kDa-cutoff Amicon centrifuge filters (Millipore), flash-frozen in liquid nitrogen and stored at -80°C. Purified protein were analysed by SDS-PAGE (NuPage Bis-Tris 4-12%, Invitrogen) with Instant Blue staining (Expedeon LTD). The final protein concentrations were determined by measuring absorbance at 280 nm and using sequence-predicted extinction coefficient and molecular weight.

### Cyclic nucleotide analysis by thin layer chromatography (TLC)

Synthesis of cyclic nucleotides by the cyclase CD-NTase was analysed by using α-^32^P labelled ATP mixed with “cold” NTPs. The reactions were carried out at 37°C for 2 hours in a buffer containing 50 mM CAPS, pH 9.4, 50 mM KCl, 10 mM MgCl_2_, 1 mM MnCl_2_, 1 mM DTT. 20 µM MkeCD-NTase were incubated with 50 µM ATP and 30 nM α-^32^P-ATP in a final volume of 20 µL. Reactions were stopped by addition of phenol:chloroform and products were isolated by chloroform extraction. Then 1 µL of the final volume was spotted on a silica gel TLC plate (Supelco Sigma-Aldrich). The plate was placed into a pre-warmed humidified chamber with running buffer composed of 30% H_2_O, 70% EtOH and 0.2 M ammonium bicarbonate, pH 9.2. Separated products were visualised by phosphor imaging. A control with a characterized Type III CRISPR system, VmeCMR ^21^ was performed with 1 µM purified VmeCMR, 2 µM target RNA incubated for 2 h at 37°C in the reaction buffer (10 mM MgCl_2_, 10 mM Tris-HCl, pH 8, 50 mM NaCl).

### Cyclic nucleotide analysis by Liquid Chromatography (LC)

To analyse the nature of the cyclic nucleotide produced by MkeCD-NTase, the previous reaction was carried out with 50 µM protein, 1 mM MnCl_2_, 250 µM ATP, 50 mM KCl, 10 mM MgCl_2_, 1 mM DTT, 50 mM CAPS, pH 9.4 and incubated 2 h at 37°C. These were diluted 2-fold with water and ultracentrifuged using 3 kDa MWCO spin filters (Pall). Liquid chromatography analysis was performed on the Dionex UltiMate 3000 system. Sample separation occurred on a Kinetiex EVO C18 2.6 µM (2.1 × 50 mm column, Phenomenex) with a 0-8% gradient of acetronitrile with 100 mM ammonium bicarbonate as the solvent. Flow rate was set at 300 µL min^-1^ and column compartment temperature was set at 40°C. Data was collected at a UV of 250 nm and a 20 µM cA_3_ commercial standard was used for comparison (Biolog).

### NADase assay

MkeTIR-SAVED NADase activity was analysed by using □-NAD^+^ as substrate – when cleaved the □-ADP_ribose_ product could be detected by fluorescence. Reactions were prepared in a 30 µL final volume with reaction buffer (50 mM Tris-HCl, pH 7.5, 50 mM KCl, 2.5 mM MgCl_2_), 1 µM cyclic trinucleotide (cA_3_) (Biolog), 0.5 µM MkeTIR-SAVED and 0.5 mM □-NAD^+^ (Sigma), or otherwise when stated in the Figure. A master mix was prepared on ice containing protein and activator, and the substrate was added immediately before beginning analysis. Reaction samples were loaded into 96-well plates (Greiner 96 half area) and fluorescence measured continuously (cycle of 20s) over 1h30 using the FluoStar Omega (BMG Labtech) with an excitation filter at 300 nm and an emission filter at 410 nm. Reactions were carried out at 28°C. A calibration curve was evaluated with value obtained after 90 min of reaction with 10, 25, 75, 225, 500 and 675 µM □-NAD^+^ as initial concentration.

### Cyclase-NADase combined assays

For the cyclase-NADase combined reaction, 5 µM MkeCD-NTase were incubated at 37°C with 250 µM ATP in 25 µL final volume containing 50 mM CAPS, pH 9.4, 50 mM KCl, 10 mM MgCl_2_, 1 mM MnCl_2_, 1 mM DTT. After 2 h, reactions were transferred into 96-well plates (Greiner 96 half area) and complemented with 0.5 µM □-NAD^+^ (Sigma). Fluorescence measurements were performed as mentioned above, during 10 min before adding 0.5 µM MkeTIR-SAVED. Reactions were carried out at 37°C for one more hour. Fluorescent data were plotted over time and the Figures were prepared with GraphPad Prism.

### Plasmid immunity assays

To analyse the effect of MkeTIR-SAVED NADase *in vivo*, we used the *Mycobacterium tuberculosis* (Mtb) Type III CRISPR system previously characterized ^20^ as an inducible producer of cA_3_ when activated by the target RNA. Here we used the following plasmids to encode MtbCsm1-5, Cas6 and a CRISPR array: (i) pCsm1-5_ΔCsm6 (*M. tuberculosis* csm1-5 under T7 and lac promoter control) and (ii) pCRISPR_TetR (CRISPR array with tetracycline resistance gene targeting spacers and *M. tuberculosis* cas6 under T7 promoter control). Competent *E. coli* C43 (DE3) cells were co-transformed with these two constructs: pCsm1-5_ΔCsm6 and pCRISPR_TetR. Plasmids were maintained by selection with 100 µg.mL^-1^ ampicillin and 50 µg.mL^-1^ spectinomycin. *Mketir-saved* gene was inserted into pRAT-Duet between NcoI and SalI restriction sites (Multiple cloning site 1). This plasmid contains the Tetracycline resistant gene targeted by MtbCsm. The plasmid immunity assay is based on the transformation of the target plasmid containing gene encoding MkeTIR-SAVED into recipient cells encoding MtbCsm. How recipient cells were prepared and transformed with the target plasmid was described previously ^20^. After the growth period in LB, cells were collected and resuspended in a LB volume adjusted to the same OD_600_ (about 0.1). A total of 3 µL of a 10-fold serial dilution were applied in duplicate to selective LB agar plates supplemented with 100 µg.mL^-1^ ampicillin, 50 µg.mL^-1^ spectinomycin, 25 µg.ml^-1^ tetracycline, 0.2% (*w/v*) D-lactose and 0.2% *(w/v)* L-arabinose. Plates were incubated overnight at 37°C. This experiment was performed with two independent biological replicates using two technical replicates for each experiment. The variant Cas10 D630A from MtbCsm was used as a control for no production of cyclic-tri-AMP.

### Analytical gel filtration

To analyse the oligomeric state of TIR-SAVED, 100 µL MkeTIR-SAVED (at least 100 µM) were injected into a size exclusion column (Superose 6 Increase 10/300 GL or Superose 12, GE Healthcare) equilibrated in 20 mM Tris-HCl, pH 8.0, 250 mM NaCl and 10% glycerol. When indicated in the Figure, 158 µM cyclic-tri-AMP (cA_3_) were added to TIR-SAVED sample before centrifugation at 10,000 *x g* for 10 min at 4°C and loaded onto the size exclusion column. Using similar gel filtration running conditions, standard proteins (#1511901, BioRad) were eluted to calculate a calibration curve of the column. 100 µL BSA (7.3 mg.mL^-1^) were injected as an additional standard.

The elution volume (V_e_) for each protein was determined based on the elution profile. Then the K_average_ (K_av_) was calculated as the ratio: K_average_ = (V_e_-V_0_)/(V_t_-V_0_) where V_0_ is the void volume (7.77 mL) and V_t_ the total volume (24 mL) of the column. The calibration curve corresponds to the K_av_ plot *versus* the molecular weight of each proteins in Log_10_. The trendline (logarithmic) was plotted with the following standard: y-globulin (158 kDa), BSA (66 kDa), ovalbumin (44 kDa) and myoglobin (17 kDa) to get the best R^2^ value (0.997) corresponding to the range of the target protein.

### Dynamic light scattering (DLS)

DLS measurements were performed with the Zetasizer Nano S90 (Malvern) instrument. In the protein dilution buffer (20 mM Tris-HCl, pH 7.5, 250 mM NaCl, 10% glycerol), 21 µM MkeTIR-SAVED were prepared with 32 µM cyclic nucleotide (cyclic-tri-AMP or others) when stated in the figure. After centrifugation at 12,000 xg for 10 min at 4°C and filtration with 0.22 µm filters, 12 µL of sample was loaded into the quartz cuvette (ZMV1012). The measurements were done at 25°C with three measures of thirteen runs. The curves of MkeTIR-SAVED WT are the mean of 3 technical replicates and 2 independent experiments.

### Small-angle X-ray scattering

SAXS datasets were recorded at the European Synchrotron Radiation Facility (ESRF, Grenoble, France) on the BioSAXS beamline BM29 ^33^ using a 2D Pilatus detector. Data were collected at room temperature (20°C) using a standard set up (automated sample mounting to a capillary by a robot)(Round et al., 2015). 100 µL TIR-SAVED (10.9 mg.mL^-1^) were injected into the Superose 12 column 10/300 GL (GE Healthcare) in 20 mM Tris-HCl, pH 8.0, 250 mM NaCl and 10% glycerol with a flow rate of 0.4 mL/min.

Sample scattering curves were obtained after subtraction of the averaged buffer signals using standard protocols with PRIMUS ^34^. The Rg and I(0) values were extracted using the Guinier approximation. Theoretical SAXS curve of the TIR-SAVED AF2 predicted structure was back-calculated and fitted with the experimental SAXS datasets with the program CRYSOL ^35^. SAXS parameters for data collection and analysis are reported in Supplementary Data Table 1.

### Electrophoretic mobility shift assays

Radiolabelled α-P^32^cA_3_ was prepared with the Type III CRISPR complex, VmeCMR, as described in the “Cyclic nucleotide analysis by thin layer chromatography (TLC)” section. The reaction product was incubated with a 3-fold dilution range of TIR-SAVED (0.07, 0.22, 0.67, 2.0, 6.0, 18 µM) for 15 min at 25°C in a final 15 µL volume. The reaction buffer was the same as for NADase activity: 50 mM Tris-HCl, pH 7.5, 50 mM KCl, 2.5 mM MgCl_2_ and 25 mM NaCl from the protein dilution buffer. Ficoll was added to the samples to get 4% as final concentration and samples were then loaded into a native 6% acrylamide gel (acrylamide:bis-acrylamide 19:1). TIR-SAVED/cA_3_ complexes were separated by electrophoresis into 1X TBE buffer, during 1 h at 200 V constant and visualized by phosphor imaging.

### Thermal shift assays

2 µM TIR-SAVED WT or variants were incubated with a range of cA_3_ concentration (0, 0.4, 2, 10 µM) in the following buffer: 20 mM Tris-HCl, pH 7.5, 250 mM NaCl, 10% glycerol supplemented by and 5X SYPRO Orange Fluorescent Dye (BioRad). A temperature gradient was applied from 25 to 95°C with 1°C incremented and fluorescence measured in Strategene MX3005. The curves are the mean of two independent experiments with technical triplicates.

### Transmission electron microscopy

Samples for screening negative-stain electron microscopy analysis were prepared by diluting purified TIR-SAVED protein alone, or with an equimolar ratio cyclic dinucleotide as indicated, to a concentration of 1 mg/ml in buffer (20 mM Tris-HCl pH 8.0, 250 mM NaCl, 10% glycerol). Electron microscopy images of negatively stained TIR-SAVED were collected using an JEOL 1200 transmission electron microscope operating at 120 keV and equipped with a Gatan Orius CCD camera at a nominal magnification of 100K ×, pixel size of 9.6 Å. 4 μl of the diluted sample was applied onto a glow-discharged 400-mesh copper grid (agar Scientific) coated with a layer of continuous carbon, followed by a 1 min absorption step and side blotting to remove bulk solution. The grid was immediately stained with 2% uranyl acetate at pH 7 and then blotted from the side and air dried prior to imaging.

Cryo-electron microscopy grids were prepared using a FEI Vitrobot Mark IV (Thermo Fisher Scientific Ltd) at 4°C and 95% humidity. 3 µl of TIR-SAVED complex was applied to Holey Carbon grids (Quantifoil Cu R1.2/1.3, 300 mesh), glow discharged for 45 s at current of 45 mA in a EMITECH K100X glow discharger. The grids were then blotted with filter paper once to remove any excess sample, and plunge-frozen in liquid ethane. All cryo-EM data presented here were collected on JEOL CRYO ARM 300 microscope, equipped with a DE36 direct detector at the Scottish Centre for Macromolecular Imaging, Glasgow, UK. 2,319 movies were collected in accurate hole centering mode using EPU software (Thermo Fisher). The CryoSPARC 3.3.1 software ^36^ was used for motion correction, CTF estimation and manual exposure correction. CryoSPARC 3.3.1 was also used for the whole single particle reconstruction workflow, from manual particle picking to classification to generate templates for autopicking and subsequent 2D classification and 3D processing, including per-particle motion correction ^37^, sharpening and 3D variability analysis ^38^ obtaining a structure with overall resolution of 3.8 Å. The final reconstruction was obtained from 596,378 particles selected from classes representing both circular and elongated particles at a 0.997 Å/pixel sampling and had overall resolution of 3.8 Å, as calculated by Fourier shell correlation at 0.143 cut-off during post-processing. The AF2 model was fitted to the map using the Chimera software, taking into account both hands. The inverted hand was the only compatible solution and was therefore chosen for further modelling and refinement using the Coot ^39^, Refmac-Servalcat as implemented in the CCP-EM suite ^40^ and PHENIX ^41^ software packages. The cryo-EM data collection, refinement and validation statistics are summarised in Supplemental Data Table 2.

## Data Availability

The electron microscopy data deposition D_1292120121 is assigned the following accession code(s): PDB ID 7QQK, EMD-14122.

**Extended Data Fig. 1.**
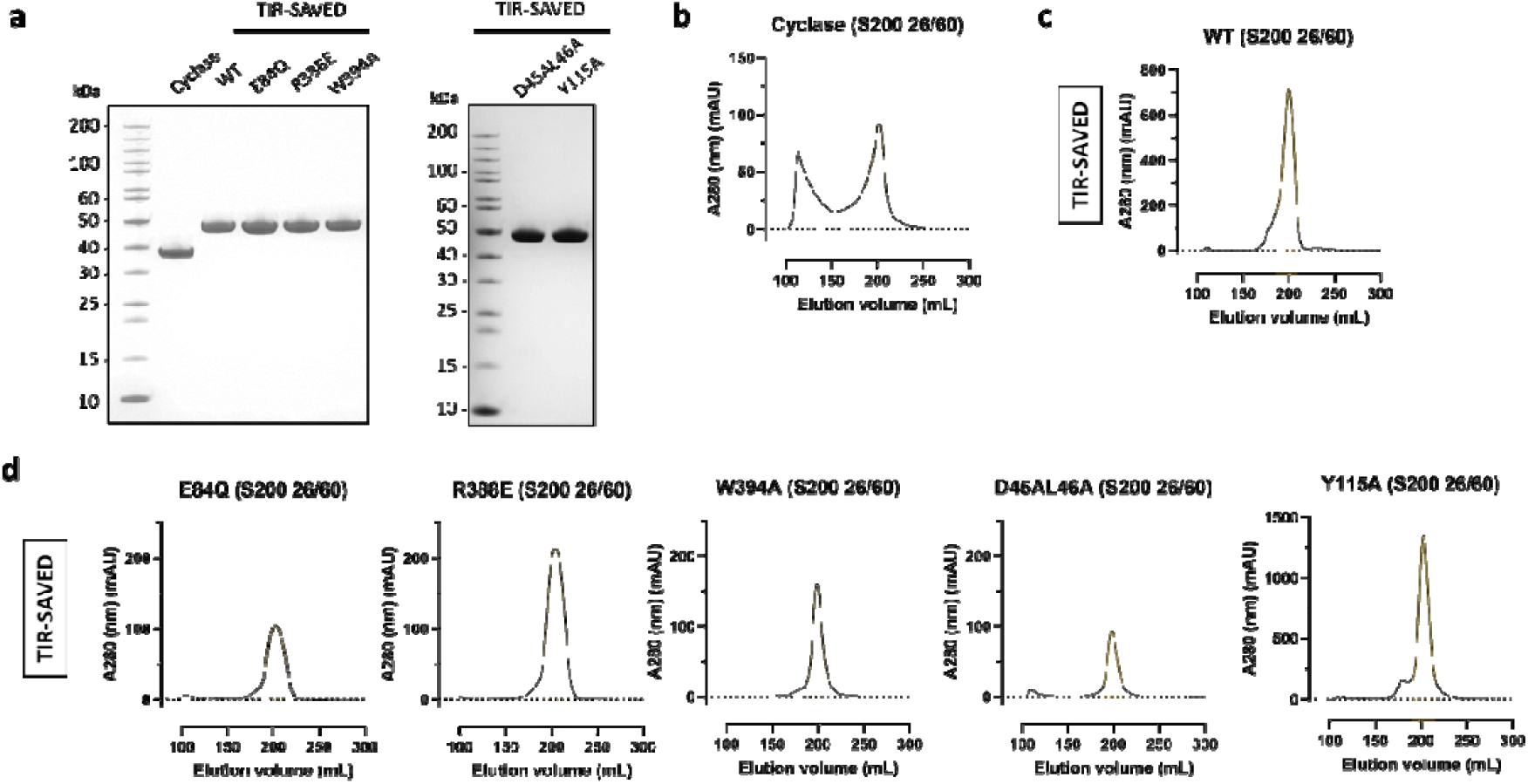
Recombinant MkeCBASS proteins. **a**, Gel electrophoretic profile of purified CD-NTase (31 kDa) and TIR-SAVED (43 kDa) proteins. Gel filtration profiles of the cyclase (**b**),TIR-SAVED WT (**c**) and TIR-SAVED variants (**d**) at the final purification step. Highlighted in orange the pool of fractions gathered and used for the further activity assays.

**Extended Data Fig. 2.**
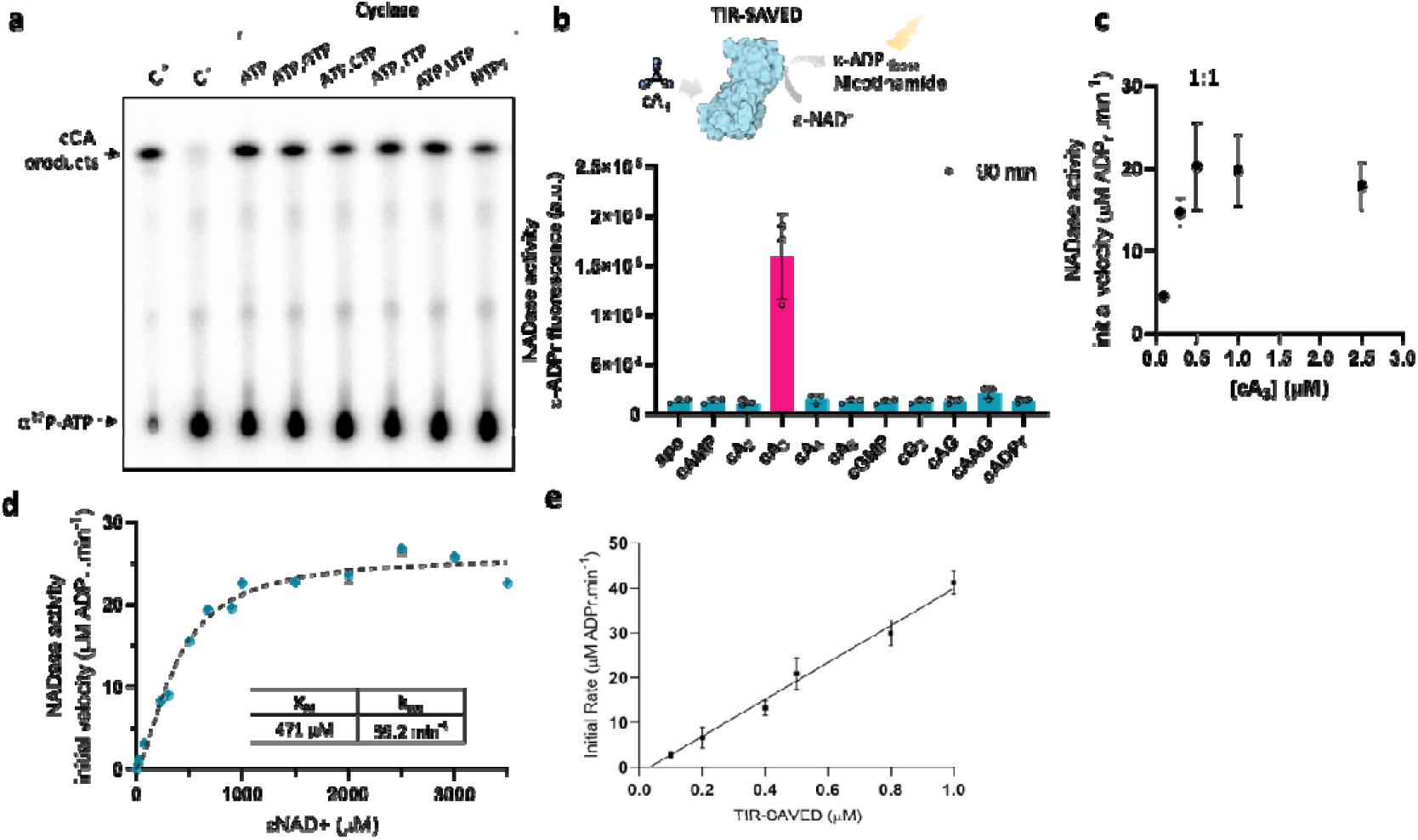
Cyclase and NADase activity. **a**, Cyclase products separated by thin layer chromatography. Radiolabelled ATP was mixed with 50 µM “cold” NTP (ATP, GTP, CTP, TTP, or UTP) and incubated with 20 µM cyclase for 2 hours at 37°C. As controls, cyclic oligoadenylate produced by the Type III CRISPR complex VmeCMR (C^+^) and the reaction without protein (C^-^). **b**, Cyclic nucleotide screening for TIR-SAVED NADase activity. 0.5 µM TIR-SAVED was incubated without (apo condition) or with 5 µM cyclic: AMP, di-AMP, tri-AMP, tetra-AMP, hexa-AMP, GMP, di-GMP, AMP-GMP, di-AMP-GMP or ADPribose. The fluorescent intensity (a.u.) at 90 min of reaction was plotted for each cyclic oligoeadenylate. **c**, Initial rate of TIR NADase activity depends on cA_3_ concentration. 0.5 µM TIR-SAVED was incubated with 0, 0.1, 0.3, 0.5, 1.0, 2.5 µM cA_3_ and 500 µM □NAD^+^. **d**, Enzymatic characterization of MkeTIR-SAVED NADase activity. Initial rate is plotted against a range of □NAD^+^ concentration. Experimental data were fitted following Michaelis-Menten equation. **e**, TIR NADase activity is proportional to TIR-SAVED concentration. 0, 0.1, 0.2, 0.4, 0.5, 0.8, 1.0 µM TIR-SAVED were incubated with 1 µM cA_3_ and 2 mM □NAD^+^. Data are the means of triplicate experiments with standard deviation (b, c, d, e).

**Extended Data Fig. 3.**
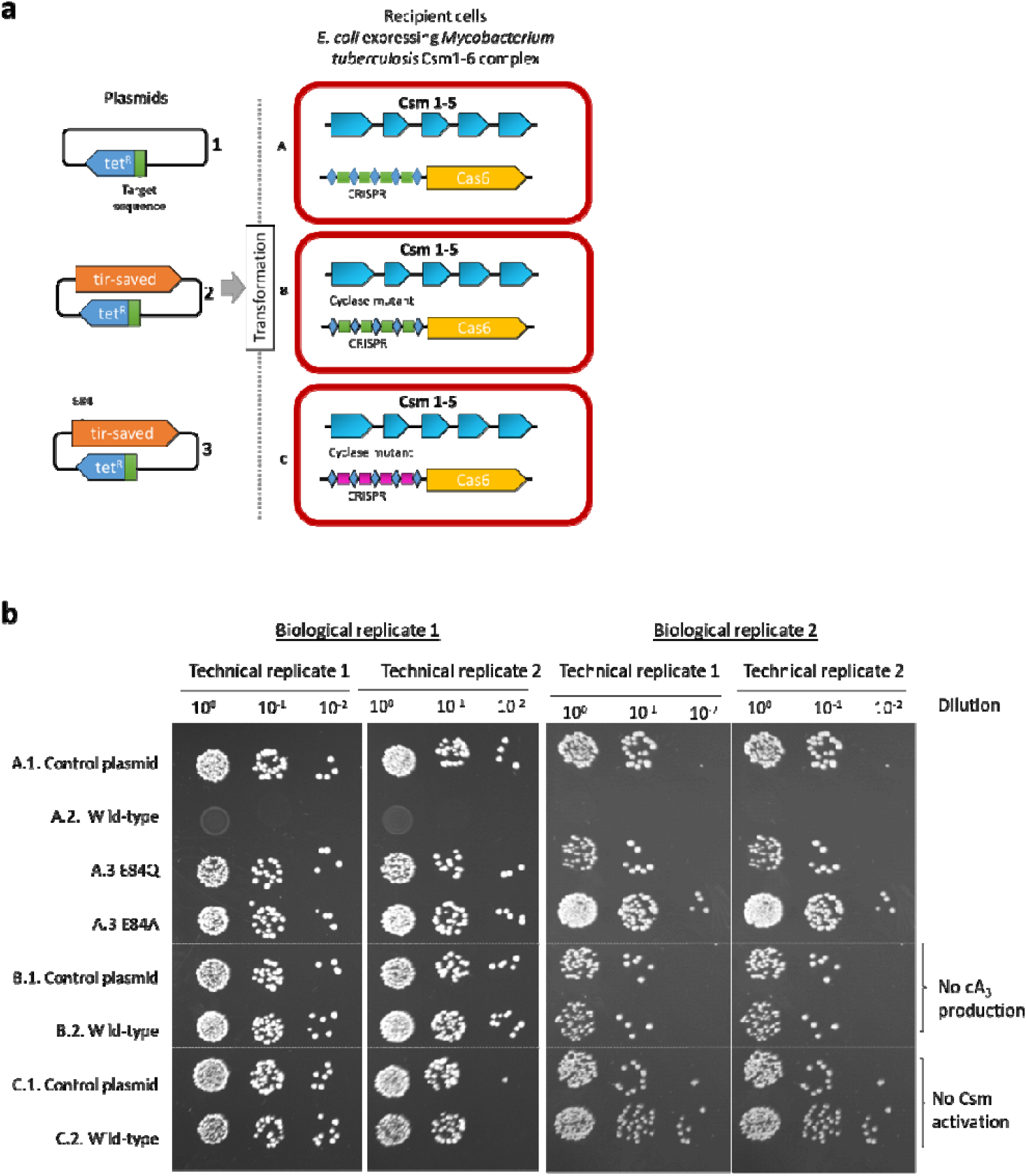
Plasmid immunity assay with MkeTIR-SAVED activated by Type III CRISPR CSM. Extended. dataset from the Fig. 1.g. **a**, Schematic of the plasmid and competent cells used for the transformation. On the left, tetracycline resistant plasmid used for to transform the recipient cell. mketir-saved gene was cloned into the multiple cloning site 1 of the plasmid 2 and 3. In the plasmid 3, the catalytic residue E84 was mutated to prevent NADase activity. Recipient cell A and B both expressed the MtbCsm targeting the tetracycline resistant plasmid. In B, MtbCsm Cas10 was mutated (D630A) to prevent cOA production and is used as a control. In recipient cell C, the tetracycline resistant plasmid is not recognized as a target by MtbCsm. **b**, Transformed colonies after incubation overnight on induced plate. The different recipient cell/plasmid combination are annotated as “A.1” (recipient cell A transformed by plasmid 1). Results from two independent experiments with technical duplicates.

**Extended Data Fig. 4.**
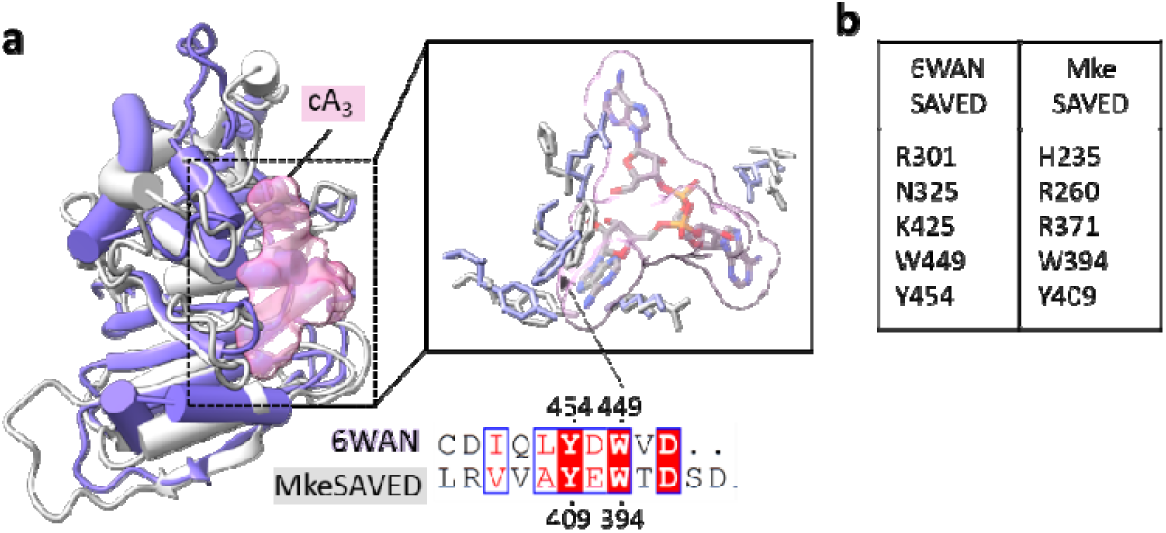
SAVED domain structure. **a**, Structural alignment of MkeSAVED domain with Cap4-SAVED cA_3_ bound structure (6WAN). **b**, Conserved residues involved in Cap4-SAVED cA_3_ binding are displayed around cA_3_ (right panel).

**Extended Data Fig. 5.**
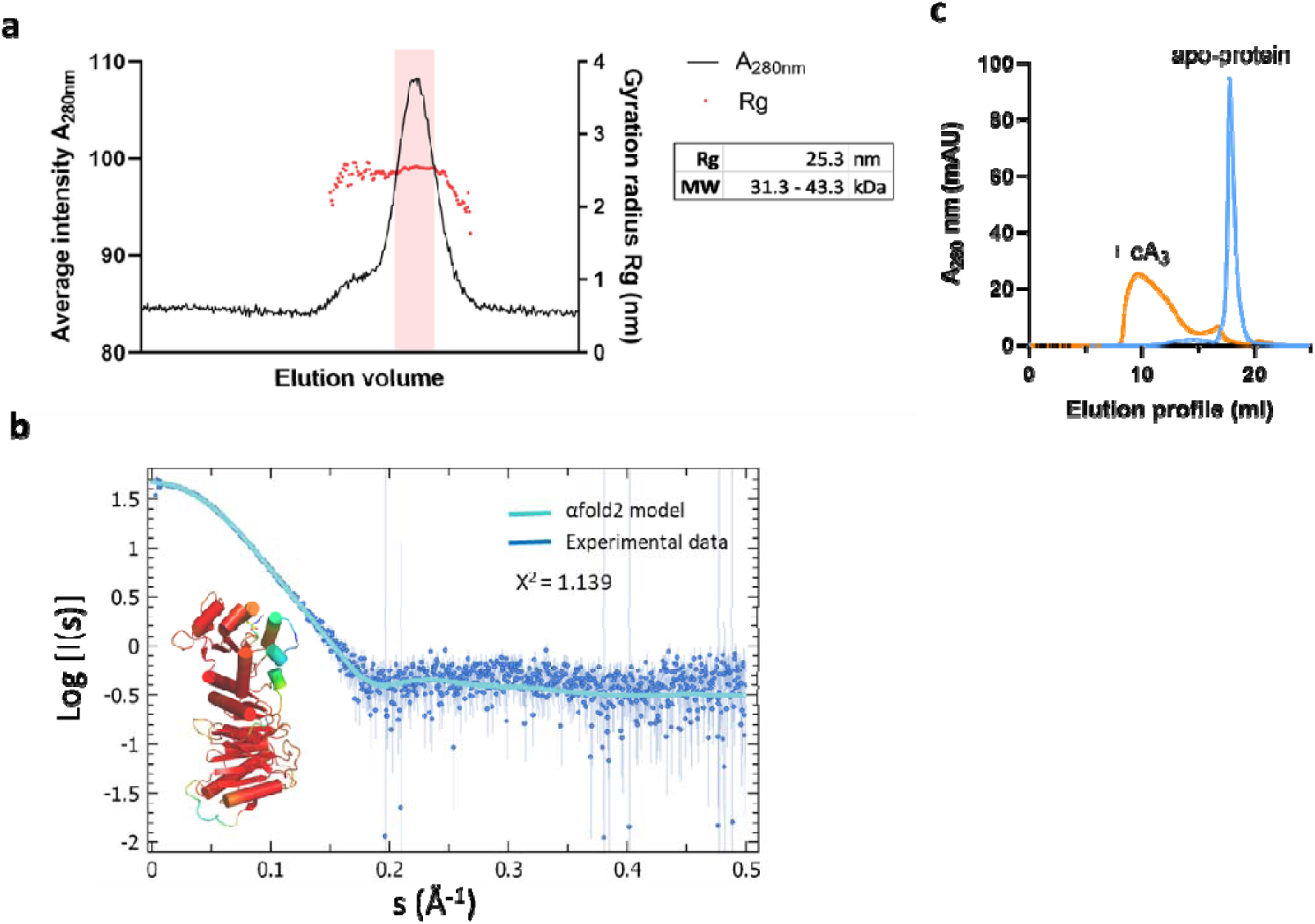
MkeTIR-SAVED is a monomeric protein in diluted solution. **a**, SEC-SAXS profile. Rg (radiation gyrus) was calculated based on Guinier approximation (see Material & Method section) and plotted against eluted volume. In red, the f ractions used to estimate the global Rg and the molecular weight range of the protein. The theoretical MW value of the recombinant MkeTIR-SAVED is 47.3 kDa. **b**, AF2 model fitted with the SEC-SAXS experimental dataset selected (red fractions in a.). **c**, Elution profile of MkeTIR-SAVED after size exclusion chromatography in absence (blue) or presence (orange) of cA_3_ for a molar ratio of 1:1.5 (protein:cA_3_).

**Extended Data Fig. 6.**
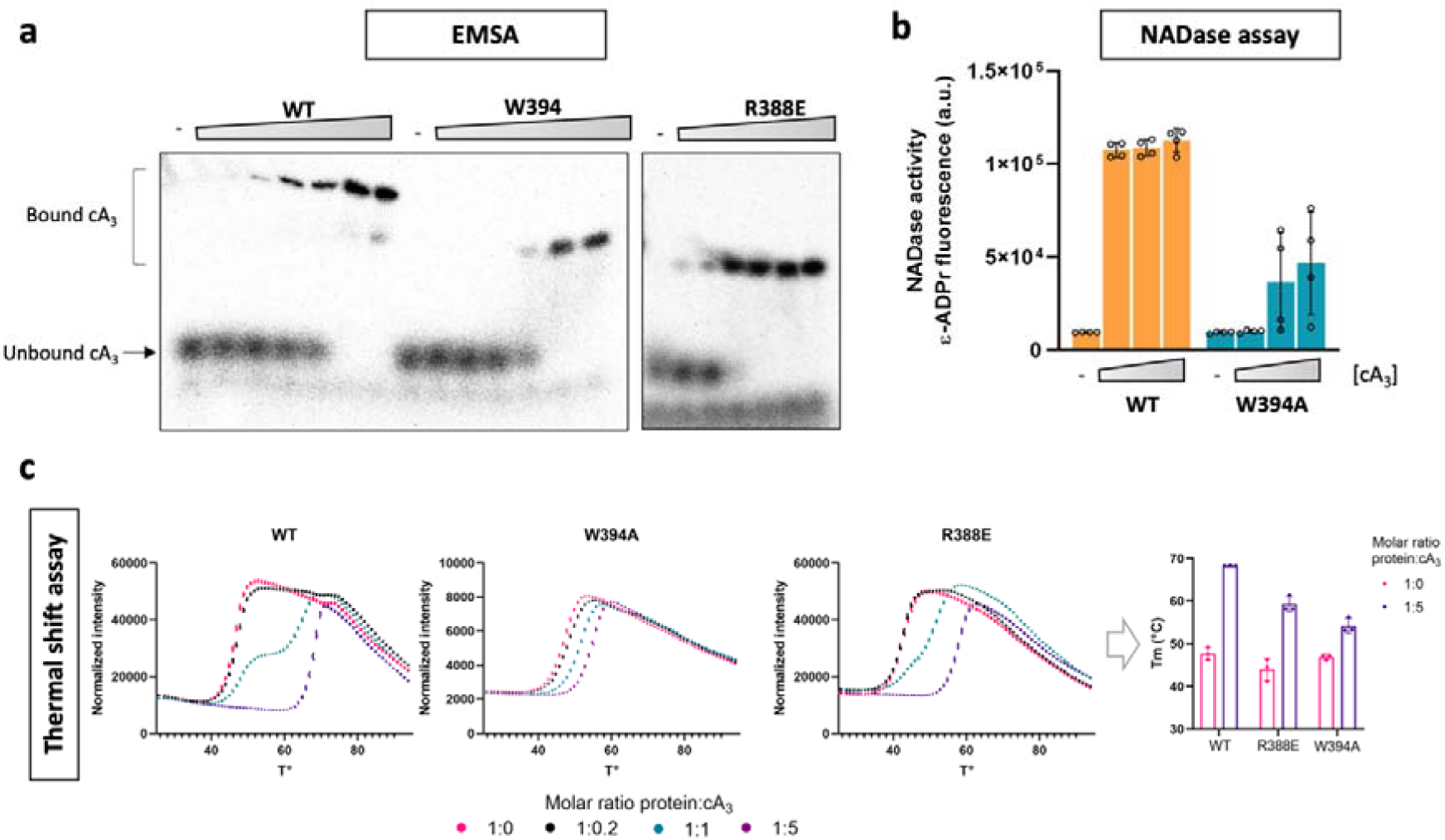
cA_3_ binding properties of TIR-SAVED mutant R388E and W394A. **a**, Electrophoretic mobility shift assay (EMSA) with radiolabelled cA_3_ incubated with a range of TIR-SAVED concentration (0, 0.07, 0.22, 0.67, 2.0, 6.0, 18 µM). Protein/cA_3_ complexes were separated by native acrylamide gel and visualised by phosphor imaging. **b**, NADase activity of W394A mutant compared to the WT TIR-SAVED enzyme. 2 µM TIR-SAVED were incubated with 0, 0.4, 2 or 10 µM cA_3_ and 500 µM □NAD^+^. The data represents the means of the fluorescent signal obtained after 60 min of reaction based on four experiments. **c**, Comparison of the thermal denaturation profile of TIR-SAVED in presence of three concentrations of cA_3_ indicated as protein:cA_3_ molar ratio of 1:0.2, 1:1, 1:5. The melting temperature Tm for the condition 1:0 and 1:5 was plotted for each protein (right panel). Experiments were done in triplicates with three measures for each temperature.

**Extended Data Fig. 7.**
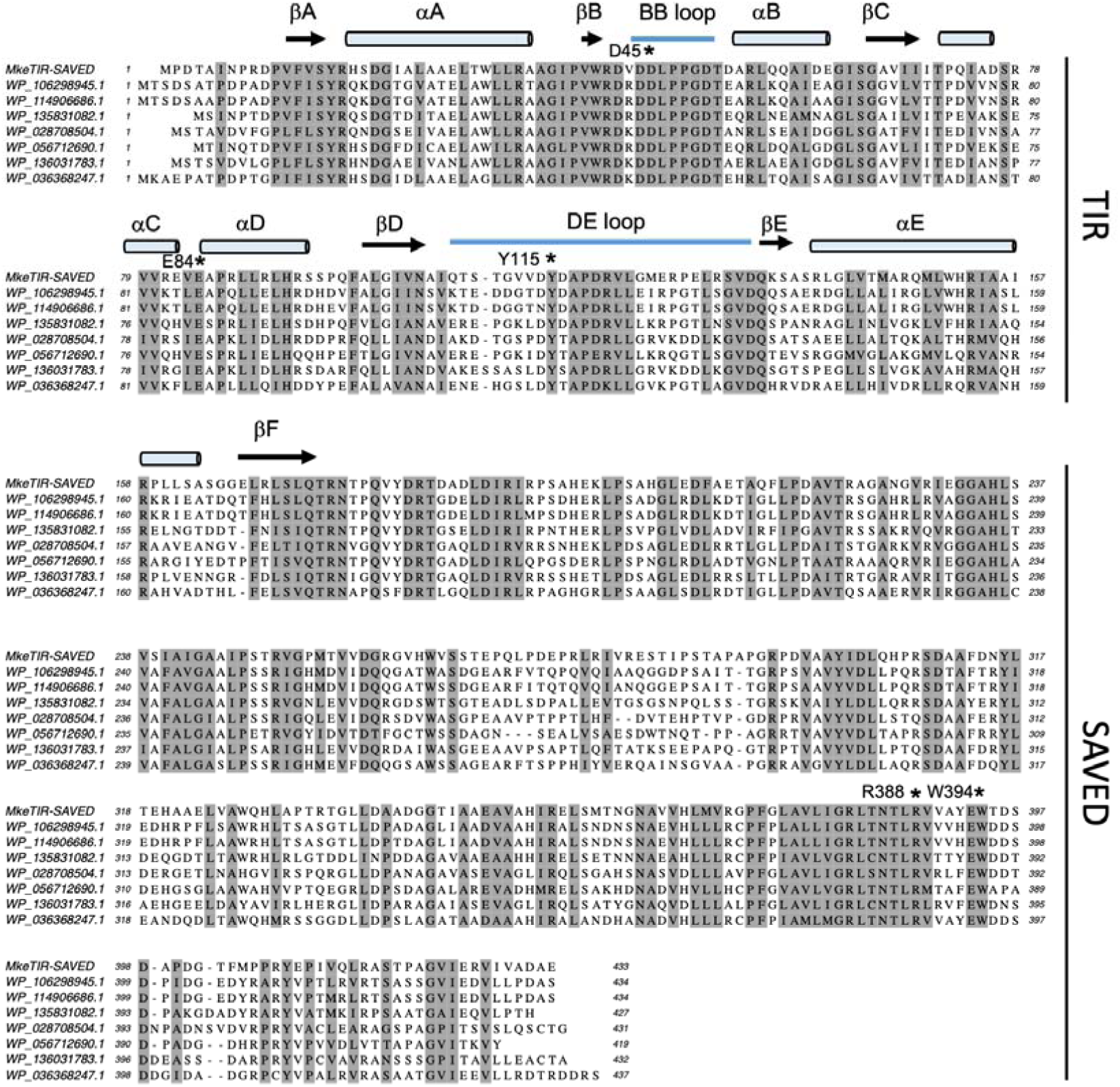
Sequence alignment of TIR-SAVED proteins. Secondary structure features are displayed for the TIR domain to match the conserved features of protein TIR family. Conserved residues are shaded. Mutated residues used in biochemical experiments are highlighted.

**Extended Data Fig. 8.**
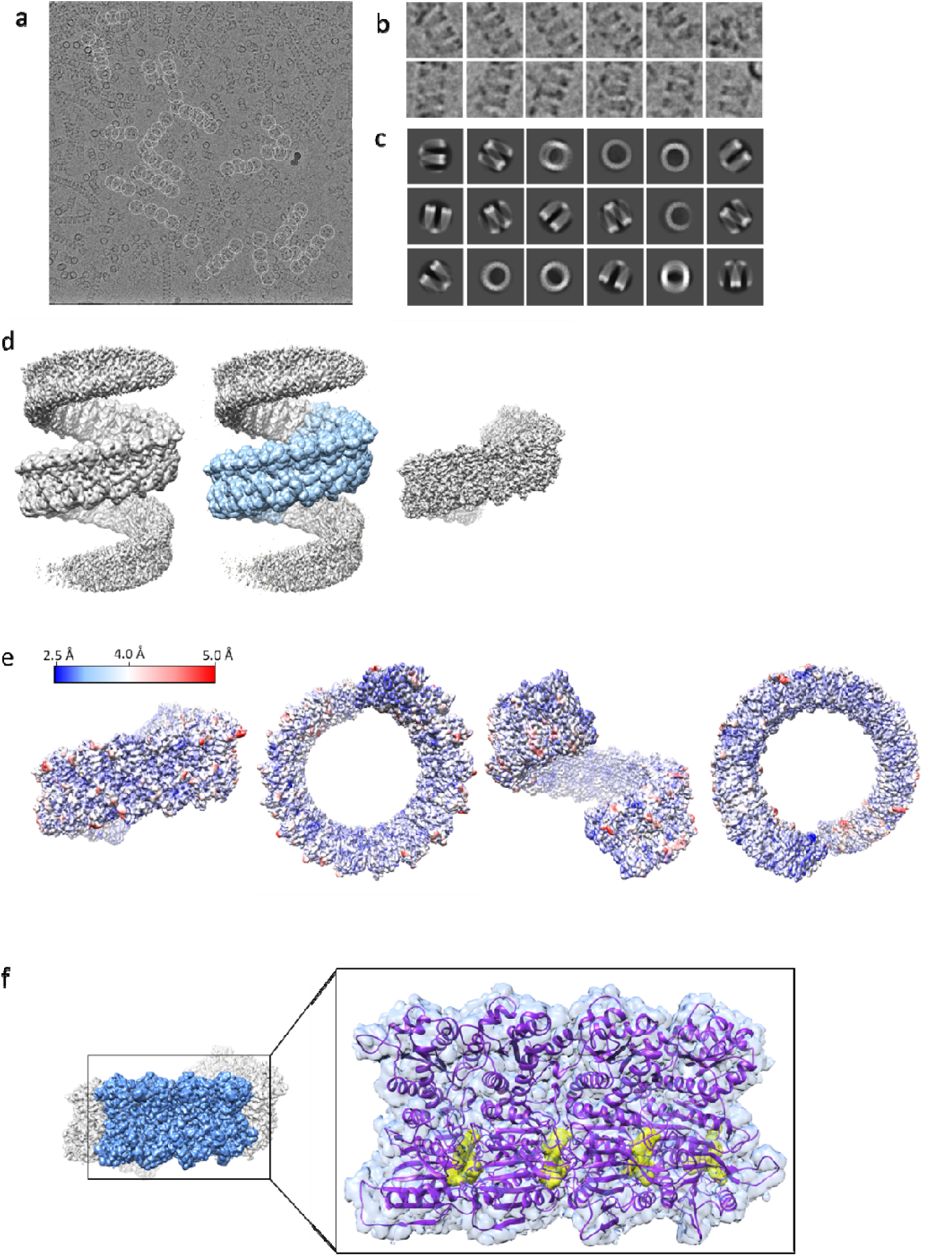
Cryo-EM analysis of the TIR-SAVED filament. **a**, Cryo-EM micrograph with single particle picked. **b**, Extract of single particle selected. **c**, 2D classes from both top and side views particles. **d**, 3D model of the filament. In blue, the extracted map used for refinement. **e**, Final refined 3D map coloured by local resolution (blue 2.5 A to red, 5 Å). **f**, Atomic model fitted into the density map for four TIR-SAVED/cA_3_ subunits.

**Extended Data Fig. 9.**
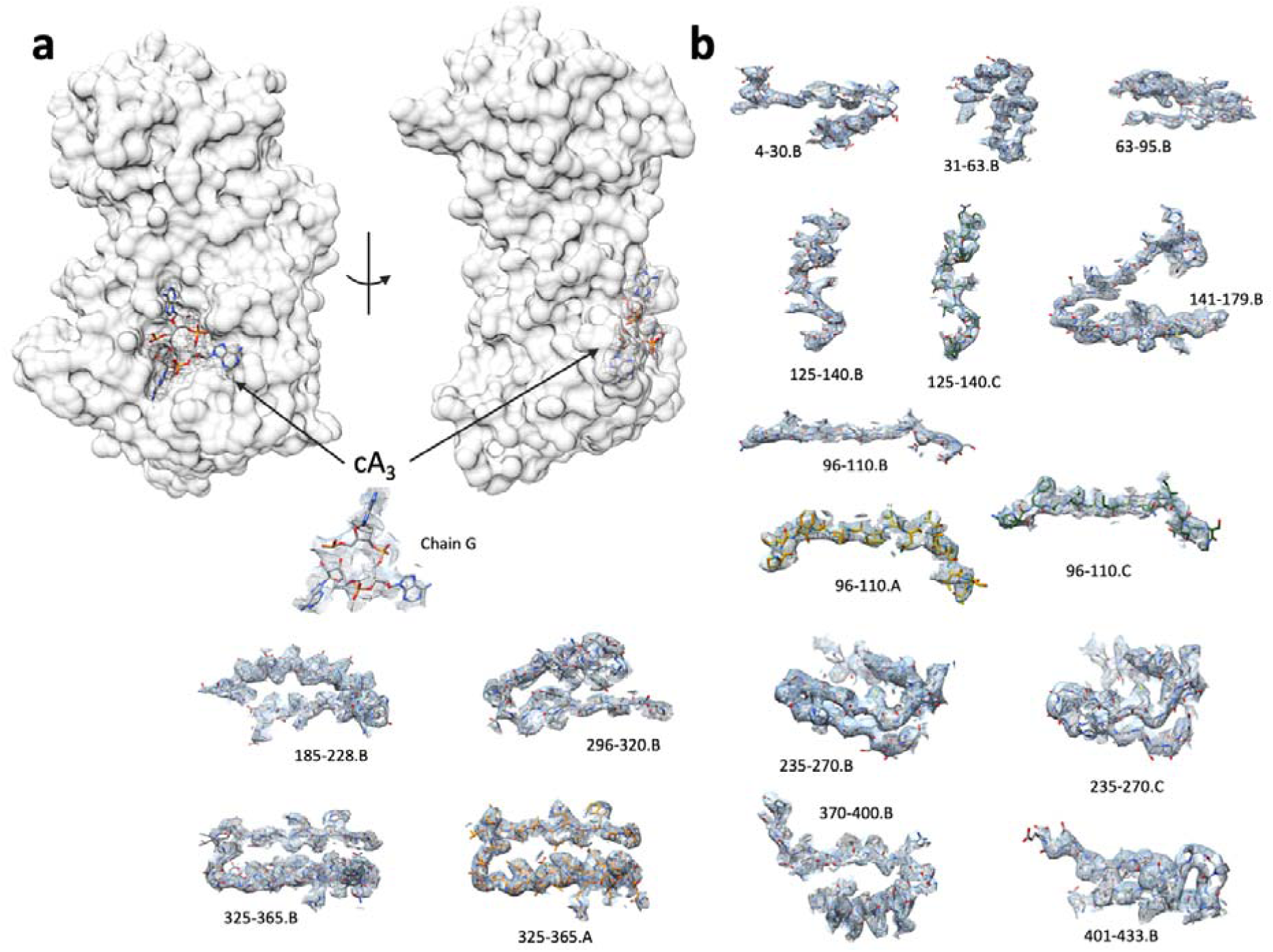
Surface representation of one TIR-SAVED subunit. **a**, position of the cA_3_ molecule bound to the SAVED protein. **b**, Representative map densities. Example map densities that allowed construction of the atomic model. The labels refer to the chain identities and residue numbers.

**Extended Data Fig. 10.**
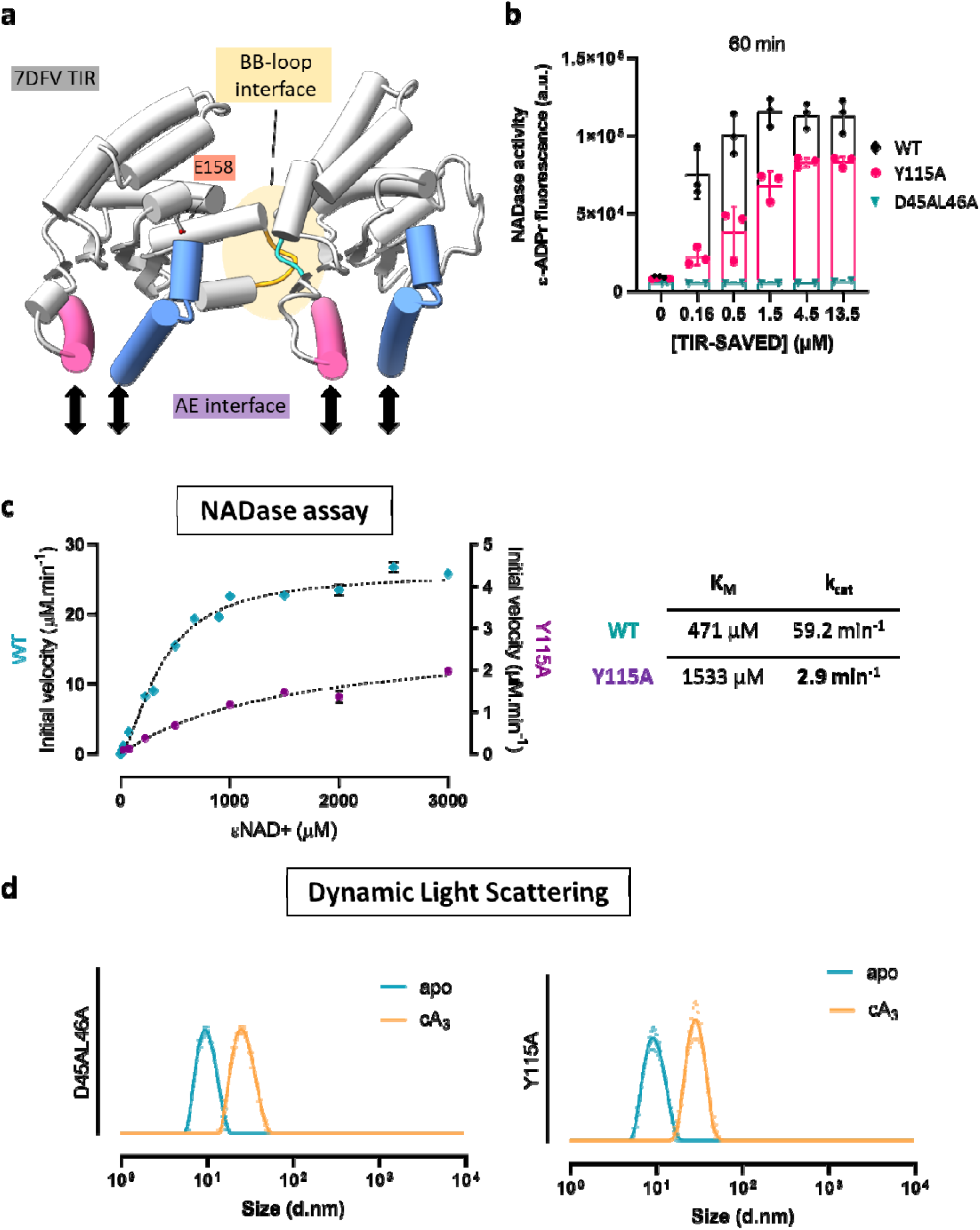
Activity and oligomerisation of TIR mutants. **a**, *Vitis rotundifolia* RUN1 TIR-TIR interfaces (PDB: 7DFV). The initial structure is organised as a tetramer of TIR-domain relying on the BB-loop and AE interaction interfaces. **b**, NADase activity comparison of the Y115A and D45AL46A mutants with the WT for a 3-fold serial dilution in protein concentrations (0.16, 0.5, 1.5, 4.5, 13.5 µM). 27 µM cA_3_ was used to activate TIR-SAVED incubated with 500 µM □NAD^+^ substrate. **c**, Enzymatic properties of Y115A mutant. Based on a NAD range concentration experiment, the initial rate of fluorescent ADP ribose production was calculated and fitted following a Michaelis-Menten model. In these experiments, as used previously for the WT, 0.5 µM Y115A were mixed in presence of 1 µM to hydrolyse 25, 75, 225, 500, 1000, 1500, 2000 and 3000 µM □NAD^+^. The right panel compares the final Michaelis-Menten parameters of Y115A with the WT protein. Data are the means of three experiments with Standard deviation error indicated. **d**, Dynamic Light Scattering analysis confirms the oligomerisation of Y115A and D45AL46A in presence of 1:1.5 protein:cA_3_ molar ratio. Experiment in technical triplicates for each condition.

## Notes

### Competing Interest Statement

The authors have declared no competing interest.

